# Mesalamine Reduces Intestinal ACE2 Expression Without Modifying SARS-CoV-2 Infection or Disease Severity in Mice

**DOI:** 10.1101/2021.07.23.453393

**Authors:** David M. Alvarado, Juhee Son, Larissa B. Thackray, Michael S. Diamond, Siyuan Ding, Matthew A. Ciorba, the IBD Investigators Group at Washington University

## Abstract

**Introduction:** Coronavirus Disease 2019 (COVID-19) is an ongoing public health crisis that has sickened or precipitated death in millions. The etiologic agent of COVID-19, Severe Acute Respiratory Syndrome Coronavirus 2 (SARS-CoV-2), infects the intestinal epithelium, and can induce GI symptoms similar to the human inflammatory bowel diseases (IBD). An international surveillance epidemiology study (SECURE-IBD) reported that the standardized mortality ratio trends higher in IBD patients (1.5-1.8) and that mesalamine/sulfasalazine therapy correlates with poor outcome. The goal of our study was to experimentally address the relationship between mesalamine and SARS-CoV-2 entry, replication, and/or pathogenesis.

**Methods:** Viral infection was performed with a chimeric vesicular stomatitis virus expressing SARS-CoV-2 spike protein and EGFP (VSV-SARS-CoV-2) and SARS-CoV-2 virus derived from an infectious cDNA clone of 2019n-CoV/USA_WA1/2020. Primary human ileal spheroids derived from healthy donors were grown as 3D spheroids or on 2D transwells. We assessed the effect of 10 mM mesalamine (Millipore Sigma) on viral RNA levels, as well as the expression of the SARS-CoV-2 receptor angiotensin II-converting enzyme 2 (ACE2), Transmembrane Serine Protease 2 (TMPRSS2), TMPRSS4, Cathepsin B (CTSB) and CTSL by qRT-PCR. 8-12 week old K18-ACE2 were treated orally with PBS or mesalamine at 200 mg/kg daily. Mice were inoculated intranasally with 1×10^3^ FFU of SARS-CoV-2. Mice were weighed daily and viral titers were determined 7 days post infection (dpi) by qRT-PCR. For the intestinal viral entry model, VSV-SARS-CoV-2 was injected into a ligated intestinal loop of anesthetized K18-ACE2 mice and tissues were harvested 6 hours post-infection.

**Results:** We found no change in viral RNA levels in human intestinal epithelial cells in response to mesalamine. Expression of *ACE2* was reduced following mesalamine treatment in enteroids, while *CTSL* expression was increased. Mice receiving mesalamine lost weight at similar rates compared to mice receiving vehicle control. Mesalamine treatment did not change viral load in the lung, heart, or intestinal tissues harvested at 7 dpi. Pretreatment with mesalamine did not modulate intestinal entry of the chimeric VSV-SARS-CoV-2 in K18-ACE2 mice.

**Conclusions:** Mesalamine did not alter viral entry, replication, or pathogenesis *in vitro* or in mouse models. Mesalamine treatment reduced expression of the viral receptor *ACE2* while concurrently increasing *CTSL* expression in human ileum organoids.

## Introduction

The coronavirus disease 2019 (COVID-19) pandemic remains a major cause of morbidity and mortality worldwide. Severe acute respiratory syndrome coronavirus 2 (SARS-CoV-2), the etiologic agent of COVID-19, is predominantly a respiratory virus but impacts many organ systems including the gastrointestinal tract.^1^ SARS-CoV-2 effectively infects and propagates in intestinal epithelial cells *in vitro* and the virus is detectable in stool and the intestinal mucosa of some patients long after clearance from the upper respiratory tract.^2^, ^3^

Some medications used to treat gastrointestinal disorders are associated with worse COVID-19 outcomes. In patients with chronic inflammatory bowel disease (IBD: Crohn’s or ulcerative colitis), corticosteroid use is associated with an increased risk of severe COVID-19.^4^ The deleterious effect of immunosuppressive corticosteroids is not unexpected and has been confirmed across non-IBD populations as well. Mesalamine (5-ASA) and its prodrug sulfasalazine have also been positively linked to increased risk of severe COVID-19 in patients with IBD^4^ and rheumatoid arthritis. This negative association is unexpected, as mesalamine has an excellent safety profile and acts primarily in the gastrointestinal tract rather than as a systemic immunosuppressant.^4^

As epidemiology studies are limited by reporting bias and unmeasured confounders, the goal of our study was to experimentally address the relationship between mesalamine and SARS-CoV-2 entry, replication, and/or pathogenesis. To address this, we used primary human intestinal epithelial cells (organoids), a mouse model of COVID-19, and a mouse model of intestinal SARS-CoV-2 entry.

## Methods

All study procedures and reagents were approved by the Washington University Institutional Review Board (#202011003) and Animal Studies Committee (Assurance #A-3381-01). Additional experimental details can be found in Supplemental Materials.

### Viruses and Cell Lines

Primary human intestinal cells were cultured as 3D spheroids and 2D monolayers as previously described.^3^ We assessed the effect of mesalamine on the expression of the SARS-CoV-2 receptor angiotensin II-converting enzyme 2 (ACE2)^5^ as well as on cellular proteases that cleave and activate the spike protein: Transmembrane Serine Protease 2 (TMPRSS2), TMPRSS4, Cathepsin B (CTSB) and CTSL.^3^, ^6^ Viral infection was performed with a chimeric vesicular stomatitis virus expressing SARS-CoV-2 spike protein and EGFP (VSV-SARS-CoV-2) and SARS-CoV-2 virus derived from an infectious cDNA clone of 2019n-CoV/USA_WA1/2020.^3^, ^7^

### Mice

8-12 week old K18-ACE2 were treated orally with PBS or mesalamine (Millipore Sigma) at 200 mg/kg daily. Mice were inoculated intranasally with 1×10^3^ FFU of SARS-CoV-2 as previously described.^7^, ^8^ Mice were weighed daily and euthanized 7 days post-infection (dpi). For the intestinal viral entry model, after three days of treatment, VSV-SARS-CoV-2 was injected into a ligated intestinal loop of anesthetized mice and tissues were harvested 6 hours post-infection (hpi).

### Statistical analysis

At least two experimental replicates were performed for all experiments. Graphs depict means and standard error of the means of biological replicates. Absolute viral copies were compared by Wilcoxon rank-sum test. Gene expression data were compared by 2-way ANOVA. All analyses were performed in RStudio.

## Results

*Mesalamine reduces* ACE2 *expression and increases* CTSL *expression in human ileal epithelial cells but does not impact SARS-CoV-2 entry*. To address the impact of mesalamine on viral entry in human intestine, primary epithelial cells were inoculated with VSV-SARS-CoV-2 with or without mesalamine. Viral entry was greater in ileum-derived epithelial cells versus colon-derived cells (**Fig. 1A**), correlating with the higher *ACE2* expression observed in the ileum rather than high *TMPRSS2* and *TMPRSS4* expression in the colon. Although mesalamine treatment reduced ileal *ACE2* expression, it did not alter VSV-SARS-CoV-2 entry. Rather, mesalamine treatment upregulated ileal expression of the lysosomal protease cathepsin L (*CTSL*), which mediates S-protein cleavage during endosomal processing of SARS-CoV-2.^6^ The reduction of ACE2 protein levels by mesalamine was confirmed in uninfected ileum enteroids by immunoblot and immunofluorescence microscopy (**Fig. 1B and C**).

**Figure 1:**
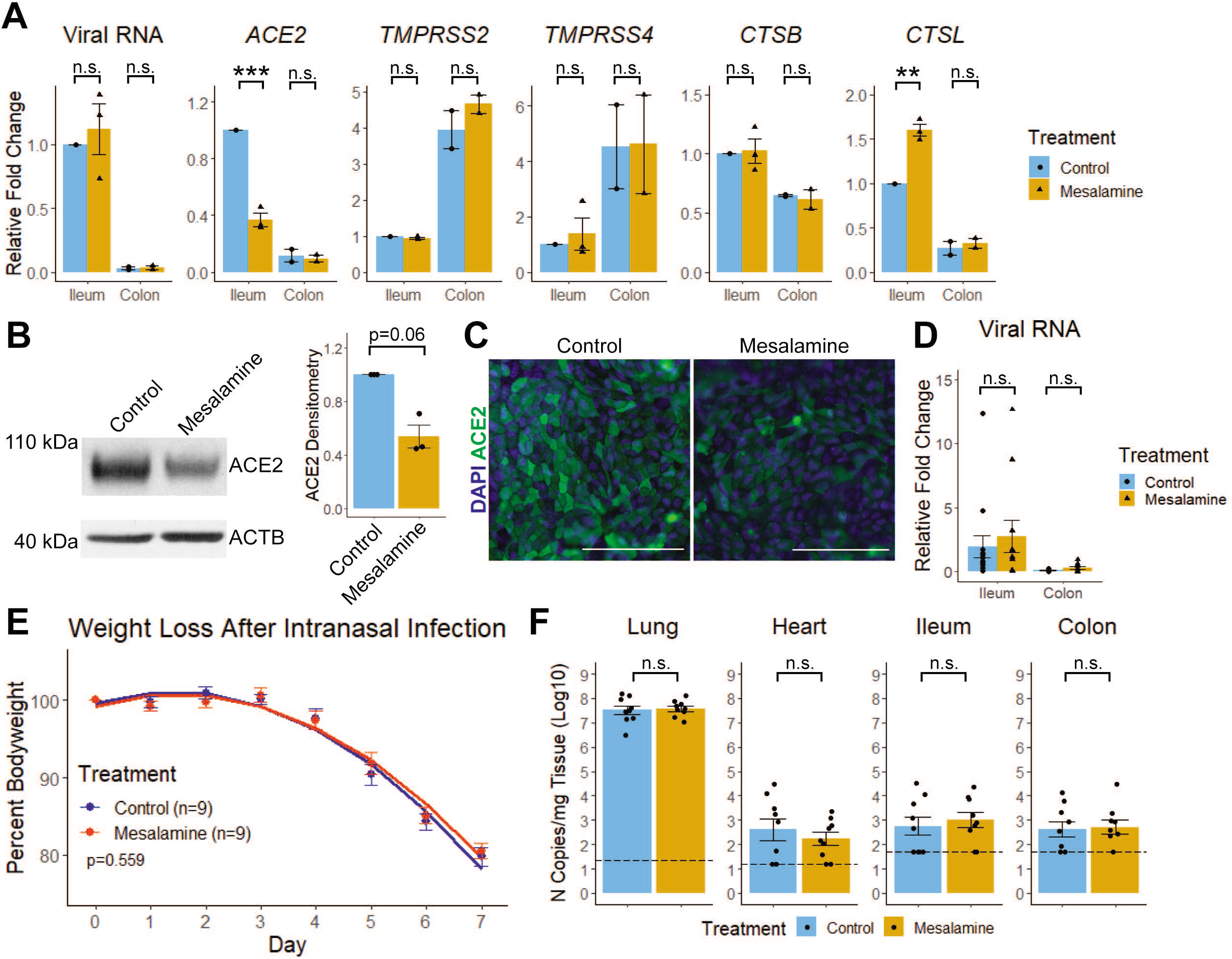
Mesalamine reduces intestinal *ACE2* expression, but does not change SARS-CoV-2 infectivity or worsen COVID-19 outcomes in mice. A) Human intestinal epithelial cells were grown as 2D monolayers and treated with 10 mM mesalamine apically 5h before administration of VSV-SARS-CoV-2 (~1.0x 10^5^ PFU/mL). After 24h, levels of viral RNA and host gene expression were determined by RT-qPCR. B) Human ileal derived epithelial cells were grown as 3D enteroids and treated with mesalamine for 30h, harvested for protein, and assessed for ACE2 by immunoblot. A representative immunoblot (left) and combined densitometry data (right) from N=3 experiments are shown. C) Apical ACE2 expression in 2D human ileal epithelial monolayers treated with vehicle (control) or 10 mM mesalamine for 30 h. Bar = 100μm. D) K18-ACE2 Tg mice were injected with VSV-SARS-CoV-2 (~1.0 x 10^5^ PFU) into the luminal space of the ileum and colon. Tissue was harvested 6 hpi and viral RNA quantification is shown. E,F) K18-ACE2 Tg mice were infected with SARS-CoV-2 (2019-nCoV/USA-WA1/2020) intranasally while receiving daily mesalamine (200 mg/kg) or sterile PBS (Control) by gavage. E) Weight loss as symptom of COVID-19. F) Viral copy quantification in tissues harvested 7 dpi. n.s., not significant; *, P<0.05; **, P<0.01; ***, P<0.001 by ANOVA.

*Mesalamine does not increase intestinal SARS-CoV-2 infectivity or morbidity in mouse models of COVID-19.* To assess the impact of mesalamine on intestinal viral entry, we developed a mouse model wherein VSV-SARS-CoV-2 is administered to ligated intestinal loops of K18-ACE2 transgenic mice (**Fig. S1A and B**). Three days of mesalamine treatment before virus administration did not alter VSV-SARS-CoV-2 levels (**Fig. 1D**) or expression of viral processing genes (**Fig. S1C**) at 6 hpi. To study the potential impact of mesalamine on SARS-CoV-2 replication and pathogenesis *in vivo*, K18-ACE2 transgenic mice were treated with mesalamine (or vehicle) three days before intranasal inoculation with 1×10^3^ FFU of SARS-CoV-2 (2019-nCoV/USA-WA1/2020).^7^, ^8^ Treatments were continued daily for 7 dpi and weight loss was monitored. Mice began losing weight 4 dpi and averaged 19% weight loss by day 7 (**Fig. 1E**). Mice receiving mesalamine lost weight equally compared to mice receiving vehicle control (p=0.559 by linear regression). Mesalamine treatment did not change viral load in the lung, heart, or intestinal tissues harvested at 7 dpi (**Fig. 1F**).

## Discussion

Mesalamine is used worldwide to treat IBD and rheumatologic disease. With early epidemiologic data associating mesalamine with poor COVID-19 outcomes,^4^ there is a need to experimentally define how this drug class interacts with SARS-CoV-2 pathogenesis. In the present study, we demonstrate that mesalamine treatment does not enhance SARS-CoV-2 infection in human intestinal epithelial cells *in vitro* or in mouse models of intestinal SARS-CoV-2. Mesalamine also does not alter viral burden or disease course in a murine model of COVID-19.

Mesalamine treatment reduced expression of the viral receptor *ACE2* while concurrently increasing *CTSL* expression in human ileum organoids. Prior studies have identified that intestinal *ACE2* expression is reduced in active ileal Crohn’s, leading the authors to posit that active IBD may be protective against SARS-CoV-2 infection.^9^ While we found that mesalamine also reduced *ACE2* expression *in vitro,* VSV-SARS-CoV-2 infection was unchanged, suggesting a potential role for additional mechanisms of viral entry.^10^ Since cathepsins have been implicated in endocytic entry of SARS-CoV-2,^6^ an area of future study will be to determine if upregulation of *CTSL* expression by mesalamine preferentially enhances spike cleavage in the endosome, thus compensating for reduced *ACE2* expression in IBD patients. Overall, these data support the safety of mesalamine use during the COVID-19 pandemic and should offer reassurance to patients with IBD and rheumatologic disorders as well as their healthcare providers.

Author names in bold designate shared co-first authorship.

## Acknowledgements

The authors thank S. Prasad, M.F. Gomez Castro, and B. Whitener for their technical support. We also thank the DDRCC AITAC for their expeditious processing of our samples, and the DDRCC PAMOC for cell culture support.

## Author contributions to manuscript

D.M.A.: study concept and design, collection and assembly of data, data analysis and integration, manuscript writing, and final approval of the manuscript. J.S.: collection and assembly of data, and final approval of the manuscript. L.B.T.: study concept and design, collection and assembly of data, and final approval of the manuscript. M.S.D.: funding, study design and scientific resources, and final approval of the manuscript. S.D.: funding, data analysis and integration, drafting, editing and final approval of the manuscript. M.A.C.: study concept and design, funding, data analysis and integration, drafting, editing and final approval of the manuscript.

All data, analytic methods, and study materials will be made available to other researchers upon request.

