## Supplemental Material for "Mesalamine Reduces Intestinal ACE2 Expression Without Modifying SARS-CoV-2 Infection or Disease Severity in Mice"

**Supplemental Methods**
No statistical methods were used to predetermine sample size. The experiments were not randomized, and investigators were not blinded to allocation during experiments and outcome assessment.

Cell Culture and Viral Preparation. VSV-SARS-CoV-2 GFP reporter chimeric virus was constructed using the SARS-CoV-2 Wuhan-Hu-1 spike and the VSV Indiana strain.[1] VSV-SARS-CoV-2 was propagated and titrated in African green monkey kidney epithelial cell line MA104 (CRL-2378.1) obtained from the American Type Culture Collection (ATCC) and cultured in complete M199 medium. SARS-CoV-2 derived from an infectious clone of 2019n-CoV/USA_WA1/2020 strain was passaged twice in Vero-TMPRSS2 cells and virus stocks were titrated on Vero-hACE2-TMPRSS2 cells. Vero-TMPRSS2 and Vero-hACE2-TMPRSS2 cells were grown in Dulbecco's Modified Eagle's Medium with 10% fetal calf serum, 10 mM HEPES pH 7.3, and 5 µg/ml blasticidin or 10 µg/ml puromycin, respectively.[2]

Primary intestinal epithelial cells were derived from biopsy of a healthy patient as previously described.[3] Briefly, biopsy was minced with scissors before digestion with dispase. Tissue was strained through a 70 µm filter and cells were embedded in Matrigel (3D culture) and maintained in 50% L-WRN conditioned medium supplemented with 10µM each Y-27632 and SB431542 as described previously. For 2D culture, transwell devices with polyester membranes with 0.4µm pore size were pre-treated with 1:40 Matrigel in PBS for 1h at 37C. Cells were trypsinized and filtered before plating on transwell membranes and cultured for ~7 days in 50% L-WRN conditioned medium with 10 µM Y-27632. Media was changed in the apical chamber every day and in the basolateral chamber every other day. Trans-epithelial electrical resistance was monitored daily and differentiation was initiated when the resistance plateaued. Differentiation was performed using Dulbecco's modified Eagle medium/F12 supplemented with 20% FBS, L-glutamine, penicillin/streptomycin, and 10 µM Y-27632 for 3 days before treatment. Mesalamine was dissolved in differentiation medium at 10 mM and pH was adjusted to ~7 with sterile 1 M NaOH. Cells were treated with mesalamine for 5 h before media was replaced with virus in the apical chamber and cells were incubated at 37C for 1 h. Viral supernatant was then replaced with fresh medium and cells were incubated at 37C for 24 h before harvest.

Animal models. Animal studies were carried out in accordance with the recommendations in the Guide for the Care and Use of Laboratory Animals of the National Institutes of Health. Virus inoculations were performed under anesthesia, and all efforts were made to minimize animal suffering. B6.Cg-Tg(K18-ACE2)2Prlmn/J mice (Jackson Laboratories #034860) were maintained in an enhanced barrier facility. Mesalamine was dissolved in sterile PBS and pH was adjusted to ~7 with sterile 1 M NaOH. Mice were gavaged daily with mesalamine or sterile PBS at a dose of 200 mg/kg starting 3 days before experiment start and continuing through termination.[4]

For the intestinal loop model, mice are anesthetized using inhaled 4% isoflurane in a chamber and maintained during the procedure by 2% by nosecone using an isoflurane vaporizer. All surgical utensils are sterilized before the procedure. The abdomen is shaved, prepped with three betadine swabs, and draped in a sterile fashion. An intestinal loop is performed according to published methods.[5] Briefly, a 3 cm midline laparotomy is performed and the intestine exposed. The terminal ileum is ligated 0.5cm proximal to the cecum with 6-0 nylon monofilament, and 100 uL of VSV-SARS-CoV-2 at ~$1x10^7$ PFU/mL is injected into the intraluminal space approximately 4cm proximal of ligature using a sterile 0.5 in 30 gauge needle for each mouse. The intestine is then ligated just distal to injection site. The proximal colon is then located and 100 uL of VSV-SARS-CoV-2 at ~$1x10^7$ PFU/mL is injected into the intraluminal space approximately 1cm distal to cecum. The proximal colon is ligated just distal to injection site to allow virus to traverse the length of the colon. Intestine is returned to the abdominal cavity and abdomen is closed in two layers starting with the abdominal wall and ending with the epidermal layer. Mice are maintained in a warmed cage until sacrifice 6h after procedure.

Intranasal infection was performed as previously described.[2] Briefly, mice were anesthetized with ketamine/xylazine cocktail (100/10 mg/kg). SARS-CoV-2 was administered to the nasal passages in 50 ul containing $1x10^3$ FFU. Bodyweight was recorded daily.

Molecular Analysis. For VSV-SARS-CoV-2 analysis, RNA was isolated from cells with Qiagen RNEasy Mini kit and from Tissue with TRIzol reagent according to manufacturers' protocols. Reverse transcriptase was performed with Applied Biosystems™ High-Capacity cDNA Reverse Transcription Kit, and real time quantitative PCR (RT-qPCR) was performed on a Bio-Rad CFX-96 system using Quantabio PerfeCTa SYBR Green Supermix. Cycle parameters were 2min at 95C and 40 cycles of 95C for 10s, 60C for 20s, and 72C for 30s. Data was analyzed by the ΔΔCq method relative to glyceraldehyde 3-dehydrogenase (GAPDH) and ileum samples of experimental controls (cells) or the average of untreated controls (animals). A list of primers used in this study is in Table 1.

SARS-CoV-2 quantification was carried out as previously described.[2] Briefly, tissues were weighed and homogenized with zirconia beads in 1 ml of DMEM media supplemented with 2% heat-inactivated FBS. Tissue homogenates were centrifuged at 10,000 rpm for 5 min. RNA was extracted using the MagMAX mirVana Total RNA Isolation Kit (Thermo Fisher Scientific) on the KingFisher Flex extraction robot (Thermo Fisher Scientific). RNA was reverse transcribed and amplified using the TaqMan RNA-to-CT 1-Step Kit (Thermo Fisher Scientific). Reverse transcription was carried out at 48 °C for 15 min followed by 2 min at 95 °C. Amplification was accomplished over 50 cycles as follows: 95 °C for 15 s and 60 °C for 1 min. Copies of SARS-CoV-2 nucleocapsid (N) gene RNA in samples were determined using a TaqMan assay. A serial dilution of RNA standard was used for copy number determination down to ten copies per reaction.

For immunoblots, 3D cultures were differentiated as for 2D cultures before treatment with 10mM mesalamine for 30h. Total protein was isolated by 1x Cell Signaling Technologies lysis buffer with protease and phosphatase inhibitors. A total of 10μg protein was resolved using a NuPAGE™ 4-12% Bis-Tris gel and protein was transferred to a nitrocellulose membrane. Antibodies used are goat anti-ACE2 (R & D AF933, 1:8,000), and rabbit anti-Actin (enQuire bio QAB10339, 1:5,000) and appropriate secondary antibodies. Densitometry was performed using Gel Analysis on ImageJ software (imageJ.NIH.gov).

For immunofluorescence, transwells were fixed in 10% formalin for 10min @RT and washed in 70% ethanol 2x5min followed by 3x5min with PBS. Goat anti-ACE2 (R & D AF933, 1:200) and AlexxaFluor 647-phalloidin (Invitrogen, 1:400) were used to visualize ACE2 and actin, respectively. Slides were mounted with hardset mounting medium with 4′,6-diamidino-2-phenylindole (DAPI, Southern Biotech). Images were collected using a Zeiss Axiovert microscope (Zeiss) and Axiocam MRM Digital black and white camera.

**Table 1. Real-time Polymerase Chain Reaction Primers Used in Current Study.**

| Target | Primer sequence 5'-3' | PrimerBank ID | Reference |
|---|---|---|---|
| VSV Nucleocapid | Fwd: GATAGTACCGGAGGATTGACGACTA | NA | 6 |
| | Rev: TCAAACCATCCGAGCCATTC | | |
| Human *GAPDH* | Fwd: AAGGTGAAGGTCGGAGTCAACG | NA | 7 |
| | Rev: TGGAAGATGGTGATGGGATTTC | | |
| Human *ACE2* | Fwd: TGAAGGCCCTCTGCACAAAT | NA | 8 |
| | Rev: ATGCTAGGGTCCAGGGTTCT | | |
| Human *TMPRSS2* | Fwd: CAAGTGCTCCAACTCTGGGAT | NA | 6 |
| | Rev: AACACACCGATTCTCGTCCTC | | |
| Human *TMPRSS4* | Fwd: CCAAGGACCGATCCACACT | NA | 6 |
| | Rev: GTGAAGTTGTCGAAACAGGCA | | |
| Human *CTSB* | Fwd: GAGCTGGTCAACTATGTCAACA | 66346650c1 | 9-11 |
| | Rev: GCTCATGTCCACGTTGTAGAAGT | | |
| Human *CTSL* | Fwd: CTTTTGCCTGGGAATTGCCTC | 125987604c1 | 9-11 |
| | Rev: CATCGCCTTCCACTTGGTC | | |
| Mouse *Gapdh* | Fwd: AGGTCGGTGTGAACGGATTTG | 6679937a1 | 9-11 |
| | Rev: TGTAGACCATGTAGTTGAGGTCA | | |
| Mouse *Ace2* | Fwd: CACTGAAGCTGGGCAGAAGT | NA | 8 |
| | Rev: TCAGCCAGTCAAACAACGGT | | |
| Mouse *Tmprss2* | Fwd: CAGTCTGAGCACATCTGTCCT | 34328226a1 | 9-11 |
| | Rev: CTCGGAGCATACTGAGGCA | | |
| Mouse *Tmprss4* | Fwd: CAACCCCTCAACAACCGTGAT | 21703806a1 | 9-11 |
| | Rev: CTCAGCAGCACTGCAATGAT | | |
| Mouse *Ctsb* | Fwd: TCCTTGATCCTTCTTTCTTGCC | 6681079a1 | 9-11 |
| | Rev: ACAGTGCCACACAGCTTCTTC | | |
| Mouse *Ctsl* | Fwd: ATCAAACCTTTAGTGCAGAGTGG | 6753558a1 | 9-11 |
| | Rev: CTGTATTCCCCGTTGTGTAGC | | |
| SARS-CoV-2 Nucleocapid | Fwd: ATGCTGCAATCGTGCTACAA | | 2 |
| | Rev: GACTGCCGCCTCTGCTC | | |
| | Probe: : /56-FAM/TCAAGGAAC/ZEN/AACATTGCCAA/3IABkFQ/ | | |

**Figure S1. Mesalamine does not change infectivity in a mouse model of intestinal VSV-SARS-CoV-2 Infection.** A) Schematic of intestinal loop model of intestinal SARS-CoV-2 infection. Wild-type and K18-hACE2 transgenic mice were gavaged daily with PBS (Ctrl) or 200mg/kg mesalamine (5-ASA) for 3 days, then VSV-SARS-CoV-2 was administered into the lumen of a ligated intestinal loop. Tissue was harvested 6h post infection for (B) immunofluorescence B) and (C) RT-qPCR. Representative data of N=3 experiments. Bars = 100µm. Graphs represent mean and SEM. n.s., not significant; **, $P<0.01$; ***, $P<0.001$ by 2-way ANOVA.

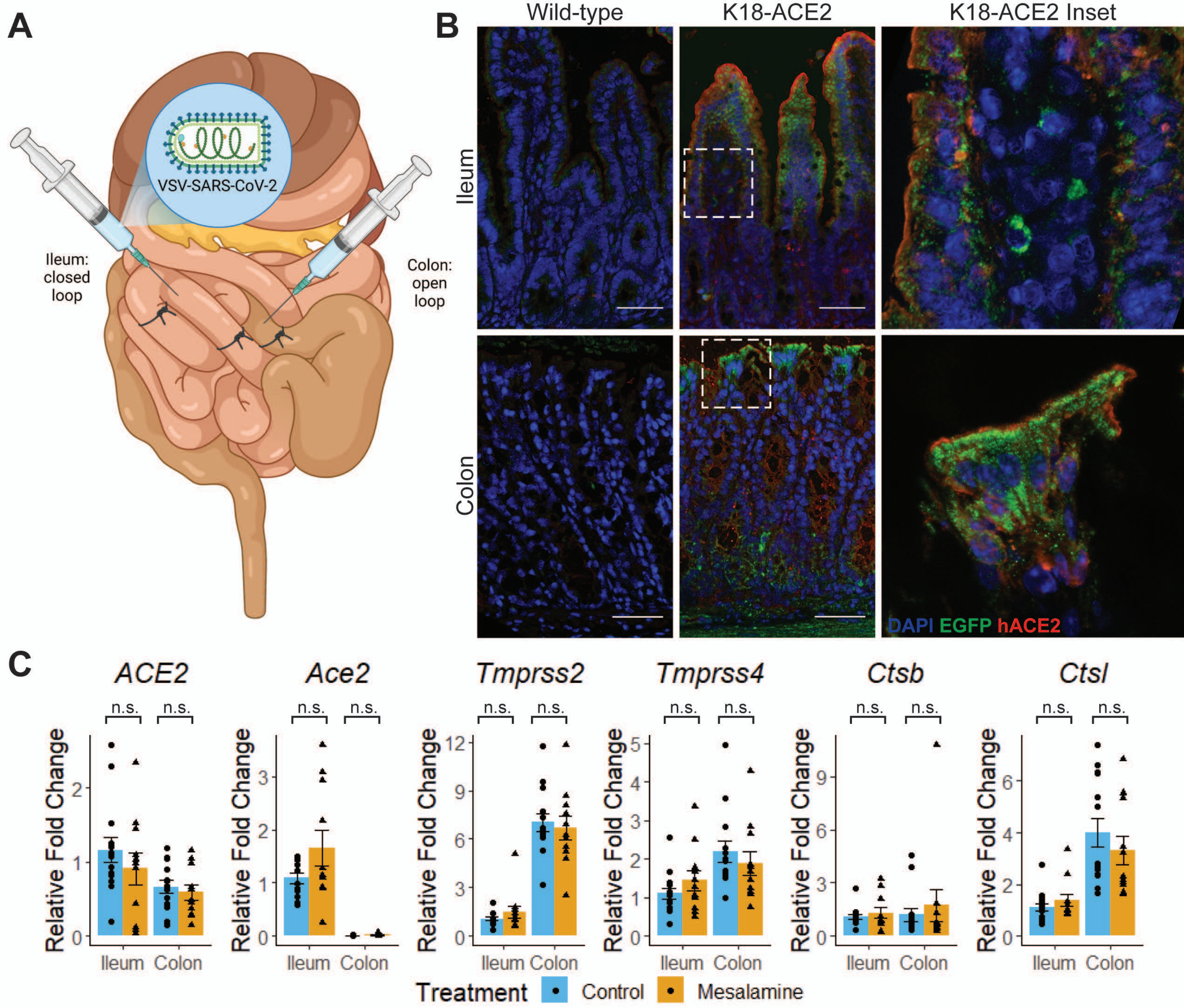
A
VSV-SARS-CoV-2
Ileum: closed loop
Colon: open loop
B
Wild-type
K18-ACE2
K18-ACE2 Inset
Ileum
Colon
DAPI EGFP hACE2
C
ACE2
Ace2
Tmprss2
Tmprss4
Ctsb
Ctsl
n.s.
Relative Fold Change
Ileum
Colon
Treatment
Control
Mesalamine
